# A congestion downstream of PSI causes the over-reduction of the electron transport chain in *pgr5* independent of membrane energization

**DOI:** 10.1101/2023.09.01.555968

**Authors:** Sandrine Kappel, Wolfram Thiele, Shany Gefen-Treves, Anita Henze, Ute Armbruster, Mark Aurel Schöttler

**Affiliations:** Max-Planck-Institut für Molekulare Pflanzenphysiologie, Am Mühlenberg 1, D-14476 Potsdam-Golm, Germany; Current address: Faculty of Biology, Rhineland Pfalz Technical University, 67663 Kaiserslautern, Germany; Molecular Photosynthesis, Heinrich-Heine-University Düsseldorf, 40225 Düsseldorf, Germany; CEPLAS – Cluster of Excellence on Plant Sciences, Heinrich Heine University Düsseldorf, 40225 Düsseldorf, Germany

## Abstract

The thylakoid protein Proton Gradient Regulation5 (PGR5) is thought to be a key component of cyclic electron flux around photosystem I. The *pgr5* mutant is characterized by impaired proton motive force (pmf) formation across the thylakoid membrane, decreased photoprotective non-photochemical quenching (NPQ), and an over-reduction of the PSI acceptor side. This over-reduction has been attributed to impaired photosynthetic control, which down-regulates plastoquinol re-oxidation at the cytochrome b_6_f complex when the lumen is strongly acidified. Here, using the *cgl160* ATP synthase assembly mutant, we show that in *cgl160 pgr5* double mutants, both the pmf across the thylakoid membrane and NPQ are fully restored to wild-type levels. However, the acceptor-side limitation of PSI in the double mutants stays comparable to the single *pgr5* mutant. This demonstrates that impaired photosynthetic control is not causal for the over-reduction of the PSI acceptor side in *pgr5*. Instead, we show that both in *pgr5* and the *clg160 pgr5* mutants, the entire high-potential chain from cytochrome f to PSI remains strongly reduced in high light. This leads to insufficient oxidizing power for plastoquinol re-oxidation by the cytochrome b_6_f complex, thus impairing pmf formation. We conclude that PGR5 plays a critical role in electron partitioning downstream of PSI.

## Introduction

In photosynthetic eukaryotes, the reactions of photosynthetic electron transport occur in the thylakoid membrane and lumen of the chloroplast. They are triggered by light-induced charge separations in the reaction centers of both photosystems (PS). Two subsequent light-induced charge separations at the chlorophyll-a dimer P_680_ in PSII lead to the reduction of plastoquinone on the stromal side of PSII. Plastoquinol dissociates from PSII and diffuses through the thylakoid membrane to neighboring cytochrome *b_6_f* complexes (b_6_f), where its re-oxidation is driven by photosystem I (PSI): A charge separation in the chlorophyll-*a* dimer in the reaction center of PSI, P_700_, leads to rapid electron transfer via the 4Fe-4S clusters on the PSI acceptor side to ferredoxin, from where electrons are predominantly used for the reduction of NADP^+^. P_700_^+^ is reduced again by plastocyanin, which in turn diffuses through the lumen to the *b_6_f* and abstracts an electron from cytochrome f. Ultimately, this provides the driving force for the oxidation of plastoquinol: One electron is directly used to reduce cytochrome f via the Rieske protein, while the second electron enters the Q cycle in a process called oxidant-induced reduction (Sarewicz et al., 2021). Usually, plastoquinol re-oxidation by the *b_6_f* is the rate-limiting step of linear electron flux, and *b_6_f* content determines the capacity of photosynthetic electron transport (reviewed by Anderson, 1992; Schöttler and Toth, 2014).

Linear electron transport is coupled to the release of protons into the thylakoid lumen by water oxidation, and by plastoquinone reduction and protonation on the stromal side of PSII and its re-oxidation at the luminal side of the *b_6_f*. The Q cycle increases the number of protons released into the lumen to three per electron, so that in total, six protons per electron pair are transported (Allen, 2002). This generates the proton-motive force (pmf) across the thylakoid membrane, which is used to drive ATP synthesis via the chloroplast ATP synthase, thereby providing chemical energy for the Calvin-Benson-Bassham cycle (CBC) and other biosynthetic pathways. Its ΔpH component also plays a major role in the feedback down-regulation of photosynthetic electron transport when the production of ATP exceeds the metabolic demand. Decreasing concentrations of orthophosphate (Pi), due to reduced consumption of ATP by the CBC itself, or due to restrictions in the metabolic use of triose-phosphates (Sharkey and Vanderveer, 1989) limit the activity of ATP synthase, so that the lumen becomes more acidic (Kanazawa and Kramer 2002; Kiirats et al. 2009; McClain et al., 2023). Lumen acidification triggers an increased dissipation of excitation energy in the PSII antenna bed via photoprotective non-photochemical quenching (NPQ). Also, lumen pH values below 6.5 slow down plastoquinol re-oxidation at the *b_6_f* in a process called photosynthetic control (Takizawa et al., 2007), which allows the short-term re-adjustment of photosynthetic electron transport to the decreased consumption of ATP and NADPH by the CBC and subsequent metabolic reactions.

It is still unclear how plants adjust to changes in the metabolic demand for ATP relative to NADPH. One possibility to selectively increase ATP production is cyclic electron flux (CEF) around photosystem I (Johnson, 2011; Nawrocki et al., 2019a). In CEF, electrons are not consumed by anabolic reactions, but re-injected into the electron transport chain. In higher plants, three pathways of CEF have been described, but their capacity and importance are still a matter of debate. Best understood is a pathway utilizing the NDH complex, a homologue of mitochondrial complex I (Laughlin et al., 2020), which functions as a proton-pumping ferredoxin-plastoquinone oxidoreductase. Per electron cycling through the NDH pathway, four protons are pumped into the lumen (Strand et al., 2017). However, in C_3_ plants, the abundance of the NDH complex usually is low (McKenzie et al., 2020). Assuming an enzymatic turnover number per complex similar to its mitochondrial and bacterial homologues, it cannot account for the high CEF rates reported in higher plants (Joliot et al., 2004; Joliot and Johnson, 2011; Nawrocki et al., 2019a). The second CEF pathway is based on direct reduction of the *b_6_f*, possibly via FNR associated with the stromal side of the complex (Zhang et al., 2001; Szymanska et al., 2011). Hardly anything is known about its capacity or regulation, but due to limited capacities of both the NDH-dependent pathway and the ferredoxin-quinone reductase (FQR) or antimycin A-sensitive pathway (see below), direct reduction of the *b_6_f* has been suggested to be the dominant pathway of CEF in C_3_ plants (Nawrocki et al., 2019a).

The third CEF pathway is the FQR or antimycin A-sensitive pathway. A first candidate component of this pathway, the Proton Gradient Regulation 5 (PGR5) protein, was identified in a screen for mutants with reduced pmf across the thylakoid membrane (Munekage et al., 2002). Apart from impaired pmf formation, the *pgr5* mutant is characterized by an over-reduction of its PSI acceptor side. This over-reduction was interpreted as a consequence of impaired NPQ and photosynthetic control in *pgr5* due to its reduced pmf formation (Munekage et al., 2002; Yamamoto and Shikanai, 2020; Zhou et al., 2023). While PGR5 does not contain any obvious cofactor-binding sequence motifs, which would be expected in a protein directly involved in electron transport, an interacting protein called PGRL1 (DalCorso et al., 2008) was shown to contain redox-active cysteine residues and speculated to bind an Fe-containing cofactor. *In vitro*, it showed activity as oxidoreductase (Hertle et al., 2013). However, inactivation of PGRL2, which is involved in the degradation of PGR5, relieves the need for PGRL1 for maintaining CEF activity, making it unlikely that PGRL1 is the oxidoreductase of the FQR pathway (Rühle et al., 2021).

Under non-fluctuating light conditions, phenotypes of the *pgr5* mutant are relatively mild, but in fluctuating light, their growth is massively impaired, and *pgr5* seedlings die within a few days. Especially PSI accumulation is strongly reduced, due to increased production of reactive oxygen species (ROS) at its acceptor side (Suorsa et al., 2012). When *pgr5* was crossed with a *sapx tapx* double mutant affected in both the stromal and thylakoid-bound ascorbate peroxidases required for ROS detoxification, the *sapx tapx pgr5* triple mutant exhibited extreme sensitivity to high light compared to its parental mutants (Kameoka et al., 2021). This was attributed to increased rates of ROS production on the PSI acceptor side. A Δ5 mutant combining knockout mutations of the oxygen evolving complex subunit genes *PSBO1*, *PSBP2*, *PSBQ1*, *PSBQ2*, and *PSBR*, suffered from a massive reduction in PSII activity, and strong growth retardation under non-fluctuating light conditions. However, when the *pgr5* mutation was introduced into the Δ5 mutant background, the sextuple mutant survived under fluctuating light conditions (Suorsa et al., 2016). The introduction of flavodiiron proteins (FLVs) from *Physcomitrella patens*, which catalyze O_2_ reduction to water using NADPH as electron donor, rescued *pgr5* under fluctuating light conditions (Yamamoto et al., 2016). Increased linear electron transport in these mutants also restored both pmf and NPQ to wild-type levels (Shikanai and Yamamoto, 2017; Yamamoto and Shikanai, 2020). Although all of these data point to a major problem on the PSI acceptor side, they were mainly interpreted as *pgr5* being deficient in proper photosynthetic control of plastoquinol re-oxidation at the *b_6_f* due to impaired CEF and reduced lumen acidification. Therefore, linear electron transport could not be properly down-regulated during high-light phases, resulting in an over-reduction of the PSI acceptor side and PSI photoinhibition due to increased ROS generation (reviewed by Shikanai and Yamamoto, 2017).

However, alternative explanations for the *pgr5* mutant phenotypes have been proposed. CEF capacity is largely unaltered in the *pgr5* mutant (Avenson et al., 2005; Nandha et al., 2007). Likewise, in the green alga *Chlamydomonas reinhardtii*, the knock-out of PGRL1 did not decrease the capacity of CEF, again arguing against a direct catalytic role of PGR5/PGRL1 in CEF (Nawrocki et al., 2019b). Avenson et al. (2005) demonstrated that in *pgr5*, ATP synthase activity was increased and suggested that a small reduction in photosynthetic ATP synthesis resulted in elevated stromal Pi and prevented a feedback inhibition of ATP synthase, thereby providing an alternative explanation for the strongly reduced pmf formation in *pgr5*. They also suggested that the restricted ATP availability limited the metabolic consumption of NADPH. In consequence, the electron transport chain was over-reduced. Somewhat similar, Tikkanen et al. (2015) and Rantala et al. (2020) concluded that PGR5 senses the redox state of PSI electron acceptors and protects PSI against photoinhibition by feedback-regulation of electron transport via photosynthetic control, possibly by downregulating ATP synthase. On the other hand, based on measurements of ferredoxin redox state in *pgr5*, Takagi and Miyake (2018) suggested a role of PGR5 in oxidizing PSI.

To distinguish between a direct role of PGR5 in pmf formation and photosynthetic control, protecting plants from over-reduction of the PSI acceptor side, and a different function of PGR5 affecting the redox state of the PSI acceptor side, we have crossed the *pgr5* mutant with the *Conserved green lineage 160* (*cgl160)* mutant. CGL160 supports ATP synthase assembly as an auxiliary protein (Rühle et al., 2014; Fristedt et al., 2015). The *cgl160* mutant suffers from moderate reduction of chloroplast ATP synthase activity, and therefore, its pmf is increased (Fristedt et al., 2015). We speculated that this reduction in ATP synthase activity might be sufficient to restore the pmf in *pgr5 cgl160* double mutants to normal levels, and might therefore largely suppress the photosynthetic defects in *pgr5*. Such a *cgl160 pgr5* double mutant was previously generated by Naranjo et al. (2021), and shown to recover NPQ to wild-type levels, but due to its strongly retarded growth, not further analyzed. Here, we show that while the pmf was indeed fully restored to wild-type levels in our *cgl160 pgr5* double mutants, and photoprotective NPQ functioned normally again, the photosynthetic electron transport chain and the PSI acceptor side remained highly reduced. Obviously, the over-reduction of the high potential chain and PSI acceptor side in *pgr5* is not caused by the absence of photosynthetic control. We conclude that PGR5 is required for correct electron partitioning downstream of PSI.

## Results

### Molecular biology, phenotypes, and plant growth

We reasoned that a defect in CEF and pmf formation in the *pgr5* mutant, which impairs photoprotective NPQ and photosynthetic control of plastoquinol re-oxidation at the *b_6_f*, could be (partly) compensated by a down-regulation of chloroplast ATP synthase, which consumes the pmf for ATP synthesis. Previously, the *Arabidopsis* CGL160 protein was shown to act as an auxiliary factor in ATP synthase assembly (Rühle et al., 2014; Fristedt et al., 2015). A reduced ATP synthase accumulation in *cgl160* resulted in decreased ATP synthase activity, increased pmf across the thylakoid membrane, and induction of NPQ and photosynthetic control already in low light (Fristedt et al., 2015). Therefore, we decided to cross the *pgr5-1* mutant with two *cgl160* mutants called *cgl160-1* and *cgl160-2* (Fristedt et al., 2015; Correa Galvis et al., 2020). Recently, additional point mutations in several photosynthesis-related genes had been observed in the *pgr5-1* mutant by whole-genome sequencing (Wada et al., 2021), but a comparison with a newly generated *pgr5*-*CAS* mutant demonstrated that these additional mutations only have a minor contribution to the reported *pgr5* phenotype (Penzler et al., 2022). To address any concerns about second side effects, we decided to also include a newly identified, independent *pgr5* mutant in our analyses, here named *pgr5-4*, with *pgr5-2* being a mutant with a single amino acid substitution (Yamamoto and Shikanai, 2019) and a third *pgr5-3* mutant being recently generated by CRISPR-Cas (Chen et al., 2023). The new *pgr5-4* mutant is in the Nossen (No-0) accession of Arabidopsis. It carries a *Ds*-transposon insertion and was obtained from Riken (Ito et al., 2002; Kuromori et al., 2004). In **Figure 1A**, the previously generated *pgr5* mutants, and the insertion site of the *Ds*-transposon in the new *pgr5-4* mutant are shown. The detailed molecular characterization of the mutant is described in the methods.

**Figure 1:**
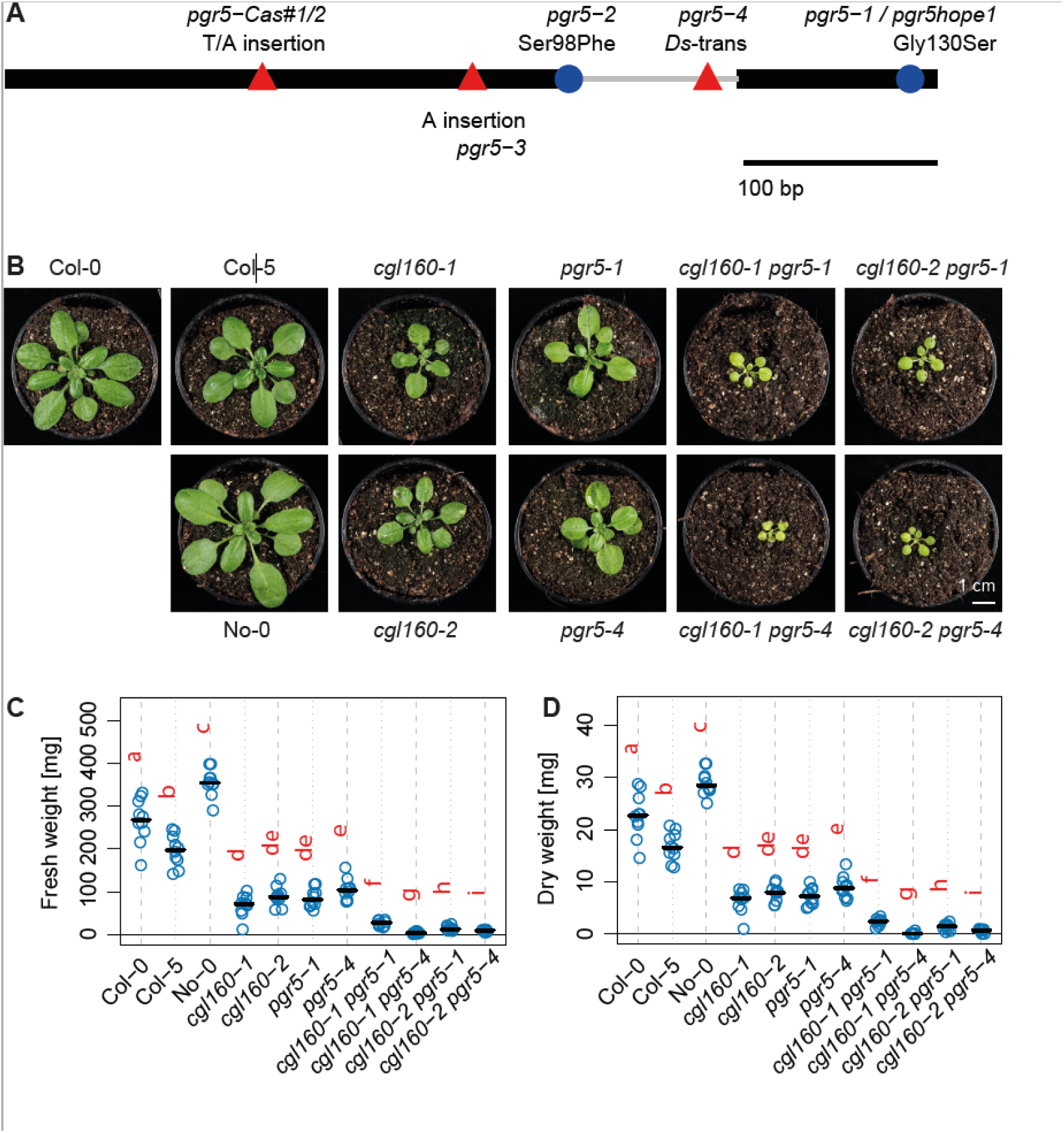
Phenotypes and growth analyses. A) PGR5 gene model with known mutations *pgr5-1* (Munekage et al., 2002), *pgr5-2* (Yamamoto and Shikanai, 2019), CRISPR Cas insertion mutants *pgr5-Cas#1* and *pgr5-Cas#2* from Penzler et al. (2022) and *pgr5-3* from Chen et al. (2023) leading to frameshifts. Coding exons are depicted as black boxes, start to stop from left to right, introns as gray lines, insertions as red triangles, substitutions as blue circles. B) Pictures of 4-week-old plants of the different genotypes. C) Fresh weights of 4-week-old plants; n= 10. Dot plot with horizontal bars representing medians. Compact letter display indicates the results of pairwise Wilcoxon Rank Sum tests, with lowercase letters representing different groups. Groups sharing the same letter are considered not statistically different (p>0.05). D) Dry weights of the plants shown in C.

As *pgr5-2* and the CRISPR-CAS9 mutant were only published after our project was initiated, we here only analyzed *pgr5-1* and our new *pgr5-4* mutant. After the four combinations of double mutants (*cgl160-1 pgr5-1*, *cgl160-1 pgr5-4*, *cgl160-2 pgr5-1* and *cgl160-2 pgr5-4*) were obtained, the three wild types, the two *cgl160* and *pgr5* single mutants, and the four double mutants were grown under non-fluctuating long-day conditions on soil. While growth of all the single mutants was somewhat slowed down in comparison to the wild types, the double mutants were massively retarded in growth (**Figure 1B**). To quantify the growth defect, fresh and dry weight of all plants was determined after four weeks of growth, shortly before the wild types started to bolt (**Figure 1C & D**). While both fresh and dry weight of the wild types differed from each other, all single mutants had clearly lower fresh and dry weight than the wild types. In all four *cgl160 pgr5* double mutants, both fresh and dry weight were again significantly reduced, relative to the single mutants. These results are similar to previously published data on a *cgl160 pgr5* double mutant by Naranjo et al. (2021). While for all subsequent analyses, the wild types were measured in the developmental state shown in **Figure 1B**, for the single and double mutants, we waited a few more days to two additional weeks, respectively, until they had reached a size closer to the wild types. All measurements were performed prior to bolting.

### Composition of the photosynthetic apparatus

To assess the composition of the photosynthetic apparatus in the *cgl160 pgr5* double mutants relative to the respective wild types and single mutants, thylakoid isolations were performed, and photosynthetic complex accumulation was assessed both by immunoblot analyses of essential subunits of the different photosynthetic complexes (**Figure 2**), and by spectroscopic quantifications of the redox-active protein components of the electron transport chain (**Table 1**). Spectroscopic quantifications of PSII, *b_6_f*, plastocyanin, and PSI were performed on a chlorophyll basis, which can serve as a proxy for the composition of the photosynthetic apparatus in the thylakoid membrane (Schöttler and Toth, 2014). Then, using the chlorophyll content per leaf area, they were re-calculated to a leaf area basis. Also, changes in the chlorophyll a/b ratio, indicative of changes in the ratio of antenna proteins (binding both chlorophyll a and chlorophyll *b*) to the reaction centers of both photosystems (binding only chlorophyll *a*), and in the maximum quantum efficiency of PSII in the dark-adapted state (F_V_/F_M_) were determined.

**Figure 2:**
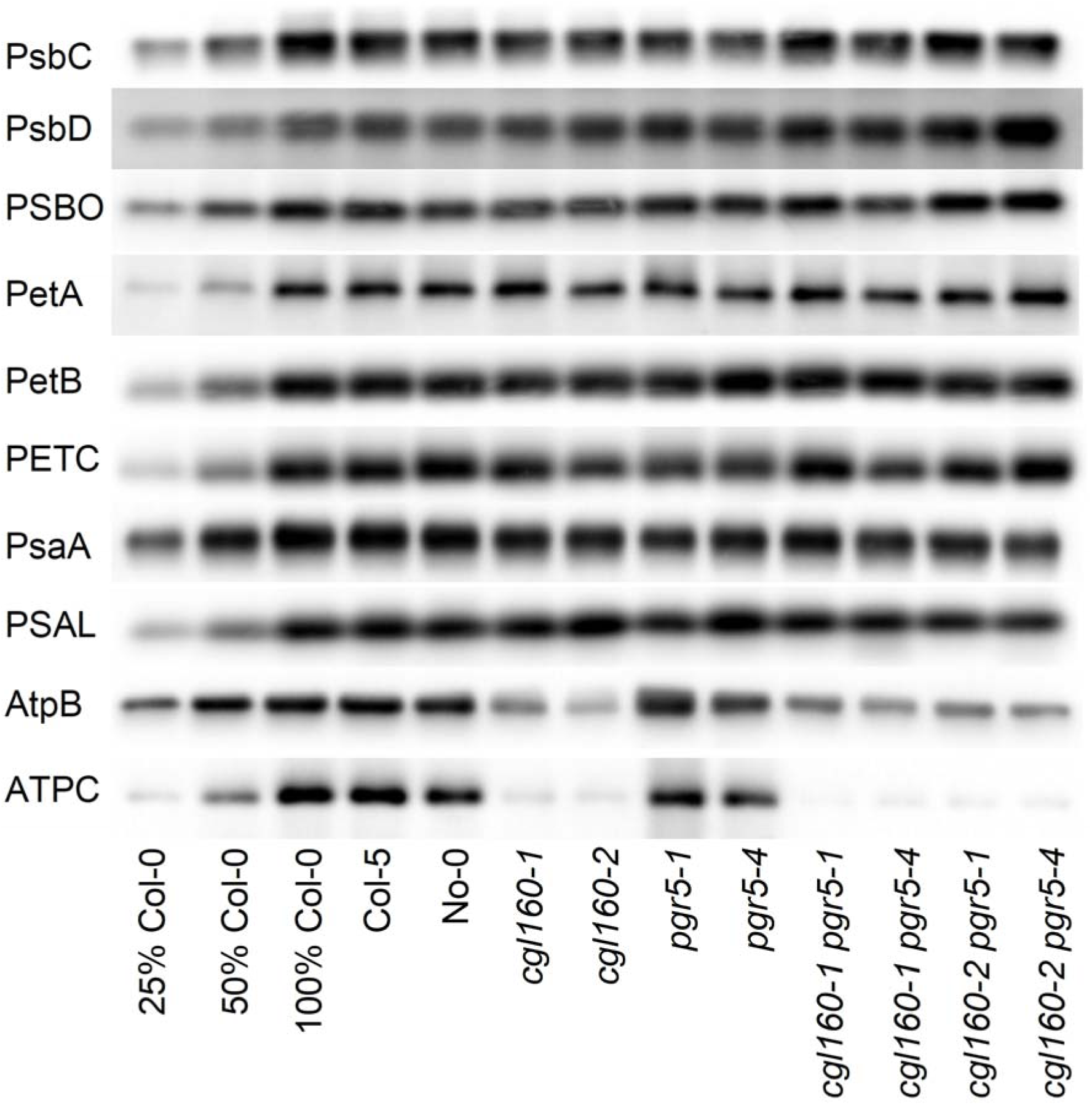
Immunoblots against subunits of PSII (PsbC, PsbD, PSBO), *b_6_f* (PetA, PetB, PETC), PSI (PsaA, PSAL), and chloroplast ATP synthase (AtpB, ATPC). Lanes one to three contain samples diluted to 25%, 50%, and a 100% sample of wild type Col-0, followed by the wild-type accessions Col-5 and No-0, the single mutants *cgl160-1* and *cgl160-2* suffering from a selective loss of chloroplast ATP synthase, the single mutants *pgr5-1* and *pgr5-4*, and the four different *cgl160 pgr5* double mutants.

While the chlorophyll content per leaf area, chlorophyll a/b ratio, F_V_/F_M_, and photosynthetic complex contents both on a chlorophyll and leaf area basis were indistinguishable between Col-0 and Col-5, the third wild type, No-0, showed differences from the other wild types in terms of reduced chlorophyll content per leaf area. While photosynthetic complex accumulation per chlorophyll was similar between all three wild types, due to the lower chlorophyll content per leaf area, most complex contents per leaf area were also slightly reduced in No-0.

In agreement with previous reports (Fristedt et al., 2015; Correa Galvis et al., 2020), the chlorophyll and photosynthetic complex contents both on a chlorophyll and leaf area basis of the two *cgl160* mutants were indistinguishable from the corresponding Col-0 wild type. The *pgr5* mutants showed minor reductions in chlorophyll content, which was slightly more pronounced in *pgr5-4*, which however is in the No-0 background with its constitutively reduced chlorophyll content. Except for a minor increase in plastocyanin content, the photosynthetic complex contents per chlorophyll were indistinguishable from the respective wild types in the *pgr5* mutants. On a leaf area basis, the accumulation of PSI was clearly reduced, relative to the respective wild type, while the accumulation of the other complexes was unaltered. Even though plants were grown under non-fluctuating light, the minor reduction in PSI may be attributable to the previously reported acceptor-side limitation of PSI in *pgr5*, possibly leading to PSI photoinhibition.

Finally, in the double mutants, a relatively large variation was observed for changes in chlorophyll content per leaf area, and photosynthetic complex accumulation. Surprisingly, this did not show any correlation with the genetic background of the crosses, in that mutants with the *pgr5-4* mutation and the No-0 background did not show more pronounced reductions in chlorophyll content and complex accumulation than mutants with *pgr5-1* and the Col-5 background. Despite the more strongly reduced biomass accumulation and impaired growth of all double mutants, their chlorophyll content per leaf area was in a similar range as or slightly higher than that of the *pgr5* single mutants. On both the chlorophyll and leaf area basis, also the contents of the major photosynthetic complexes of the double mutants were very similar or even higher than in the single *pgr5* mutants. Therefore, chlorophyll content and photosynthetic complex accumulation do not provide any clear explanations for the reduced growth of the *cgl160 pgr5* double mutants.

For immunoblots, equal amounts of chlorophyll were loaded per lane (**Figure 2**). The accumulation of essential subunits of the major photosynthetic complexes was tested. For semiquantitative analyses, a dilution series of the Col-0 wild type to 25%, 50% and 100% of the sample was used. In agreement with the spectroscopic complex quantifications, the immunoblots against the chloroplast-encoded PSII reaction center subunits **PsbD** (the D2 protein) and **PsbC** (the inner antenna protein CP43) revealed no major differences in PSII content in the different wild types and all mutants. Likewise, accumulation of the nuclear-encoded oxygen-evolving complex subunit **PSBO** was unaltered. In case of the *b_6_f*, no clear differences between wild types and mutants were observed for the accumulation of the plastome-encoded **PetA** (cytochrome f) and **PetB** (cytochrome b_6_) subunits and the nuclear-encoded **PETC** subunit (the Rieske protein). The same was true for PSI, for which antibodies against the chloroplast-encoded **PsaA** reaction center core subunit, and against the nuclear-encoded **PSAL** subunit were used. Finally, accumulation of ATP synthase was tested using antibodies against the chloroplast-encoded CF1 subunit **AtpB**, which forms part of the catalytic center of the ATP synthase, and against the nuclear-encoded **ATPC** protein. As expected, these immunoblots revealed drastic reductions in ATP synthase content to less than 25% of wild-type levels both in the two *cgl160* single mutants and in the four *cgl160 pgr5* double mutants, in agreement with previous reports (Rühle et al., 2014; Fristedt et al., 2015; Correa Galvis et al., 2020). These blots also show that ATP synthase content is not altered in the *pgr5* single mutants. Clearly, plants do not compensate for impaired pmf formation in the *pgr5* mutant by adjusting ATP synthase content.

### In the *cgl160 pgr5* double mutants, NPQ is fully restored, but linear electron transport is severely impaired

To understand the basis of the severe growth retardation of the double mutants, we performed an in-depth characterization of their photosynthetic performance, starting with light response curves of the chlorophyll-*a* fluorescence parameters linear electron transport (ETRII), non-photochemical quenching (NPQ), and Y(NO), a measure for non-regulated dissipation of excitation energy in PSII, which points to an increased propensity to PSII photoinhibition. First, we compared light response curves of the three different wild types to the two single mutants of *pgr5* and *cgl160*, respectively (**Figure 3**, left column). Because light response curves for all three parameters were basically identical for the three wild types and between the two *cgl160* mutants, only the curves for Col-0 and *cgl160-1* were re-used in the right column, where they were compared to the curves of the four *cgl160 pgr5* double mutants. In case of the two *pgr5* mutants, the left column revealed minor differences between *pgr5-1* and *pgr5-4* for all chlorophyll-*a* fluorescence parameters, with *pgr5-1* being slightly more affected than *pgr5-4*. Despite these small differences, for clarity of presentation, in the right column, only the light response curve of *pgr5-1* was re-used.

**Figure 3:**
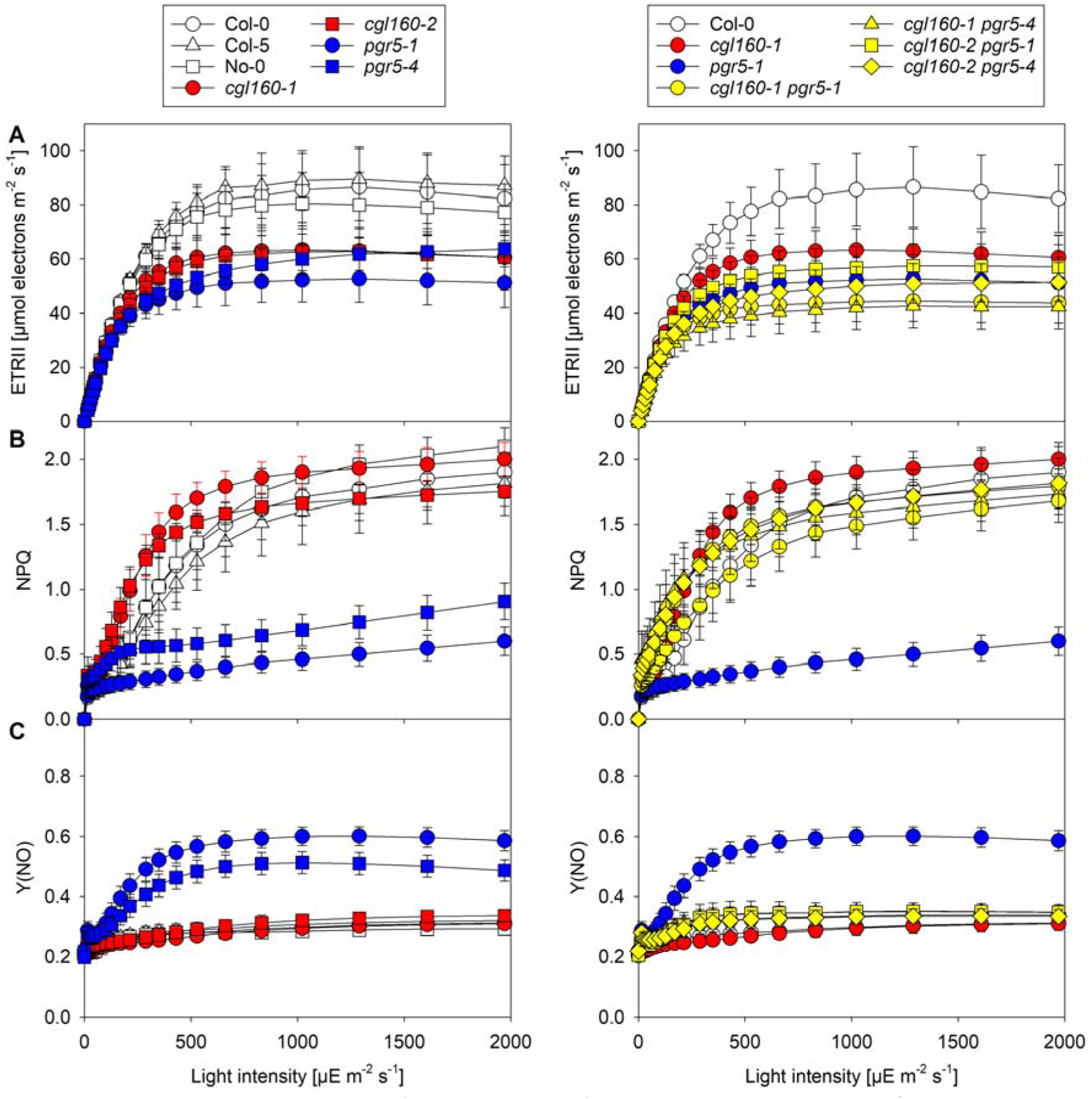
Light response curves of chlorophyll-*a* fluorescence parameters. A) Linear electron transport (ETRII). B) Non-photochemical quenching (NPQ). C) Y(NO), a measure for the non-regulated dissipation of excitation energy in PSII. Left column: Data for the three wild-type accessions and the two single mutants of *cgl160* and *pgr5*, respectively. Right column: Data for Col-0, *cgl160-1*, *pgr5-1* and for the *cgl160 pgr5* mutants.

Light response curves of linear electron transport confirmed that in all single mutants, linear electron transport was reduced by up to 50%, relative to the three wild types accessions. This is in line with previous reports (Fristedt et al., 2015; Correa Galvis et al., 2020). Furthermore, in line with the strongly retarded growth of the *cgl160 pgr5* double mutants, their linear electron transport capacity was also strongly decreased (**Figure 3A**). As expected, in both *pgr5* single mutants, NPQ induction was strongly impaired (**Figure 3B)**, while in the *cgl160* mutants, NPQ induction was slightly shifted to lower light intensities, as previously shown (Fristedt et al., 2015). In all *cgl160 pgr5* double mutants, NPQ was fully restored to wild-type levels or even induced at slightly lower light intensities, similar to the *cgl160* mutants (**Figure 3B**). While Y(NO) was indistinguishable in the wild types and *cgl160* mutants, both *pgr5* mutant lines showed a clearly increased Y(NO) in higher light intensities. Because their NPQ induction is compromised, more light is dissipated in a non-regulated manner in these mutants, potentially rendering them more sensitive to photoinhibition. Y(NO) in all *cgl160 pgr5* double mutants resembled the wild types, in line with their restored NPQ (**Figure 3C**).

### ATP synthase activity and pmf

The restored NPQ in the *cgl160 pgr5* double mutants suggested that also thylakoid membrane energization was fully functional again. To directly confirm pmf restoration, light response curves of the electrochromic shift (ECS), a measure for the light-induced pmf across the thylakoid membrane were determined (**Figure 4A**). To this end, steady-state illumination at the selected actinic light intensities was interrupted by short intervals of darkness, during which the pmf rapidly collapsed, due to rapid proton efflux through the ATP synthase. The maximum amplitude of the ECS, ECS_T_, was then used as a measure for the light-induced pmf. For all plants, ECS_T_ was measured at 1295, 311, 148 (equivalent to the growth light intensity), and at 88µE m^-2^ s^-1^. In case of Col-0, and additional light intensity of 193 µE m^-2^ s^-1^ was measured. In all lines, ECS_T_ increased strongly up to a light intensity of 311 µE m^-2^ s^-1^, approximately twice the growth light intensity, and then showed only a moderate further increase with light intensity (**Figure 4A**). As expected, in the *cgl160* mutants, ECS_T_ was increased both under light-limited and light-saturated conditions, in line with their induction of NPQ at lower light intensities. On the other hand, ECS_T_ was clearly reduced in both *pgr5* mutants, also in agreement with their impaired NPQ. In line with the full recovery of NPQ, also thylakoid membrane energization of the *cgl160 pgr5* double mutants was as high or even slightly higher than in the wild types.

**Figure 4:**
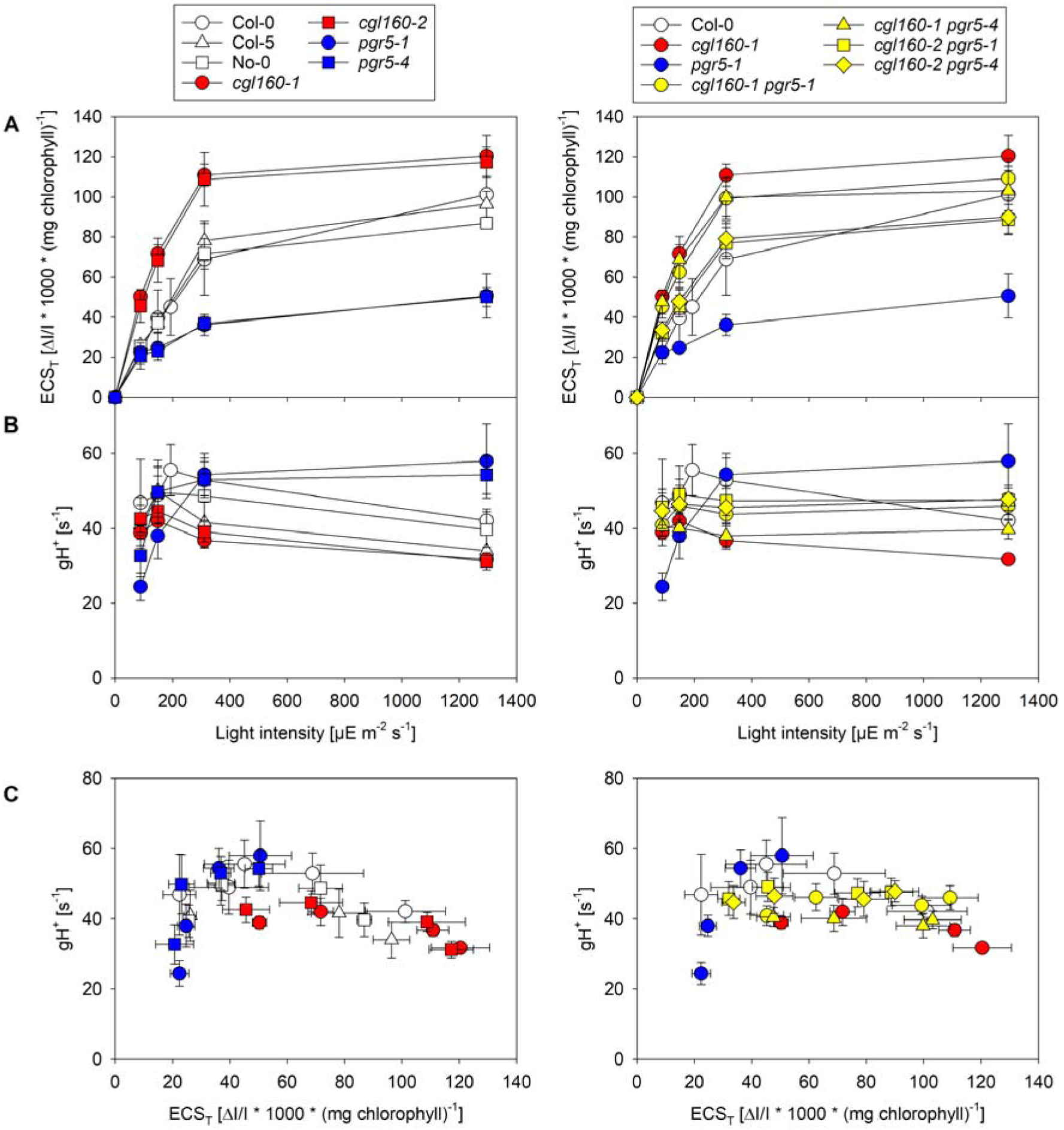
Light response curves of (A) ECS_T_ and (B) gH^+^, and blot of ECS_T_ versus gH^+^ for the three wild-type accessions and the *pgr5* mutants (C). Left column: Data for the three wild-type accessions and the two single mutants of *cgl160* and *pgr5*, respectively. Right column: Data for Col-0, *cgl160-1*, *pgr5-1* and for the *cgl160 pgr5* mutants. Shown are average values and standard deviations for n = 9 replicates for all wild types and mutants.

Next, we fitted the ECS dark interval relaxation kinetics with a single exponential decay function and used the reciprocal value of the time constant as a measure for the thylakoid conductivity for protons (gH^+^) and therefore for ATP synthase activity (**Figure 4B**). At 1295 µE m^-2^ s^-1^, we observed the highest ATP synthase activity for both *pgr5* mutants (as described by Avenson et al., 2005), while for all other mutants, ATP synthase activity was partly inhibited, relative to the gH^+^ values measured at 311 and 148 µE m^-2^ s^-1^. Over the entire light intensity range, ATP synthase activity in both *cgl160* mutants was clearly decreased, relative to their Col-0 wild type. However, this reduction was much less pronounced than that in ATP synthase content (**Figure 2**). It had been previously shown that an up to 50% reduction in ATP synthase content does not affect its activity, while further reductions result in proportional decreases in catalytic activity (Fristedt et al., 2015; Rott et al., 2011; Strand et al., 2023). The *cgl160 pgr5* double mutants showed ATP synthase activities between the wild type and the *cgl160* single mutants.

Remarkably, when ATP synthase activity in the wild types increased at decreasing actinic light intensity, its gH^+^ became indistinguishable from the two *pgr5* mutants (**Figure 4B**). To test if this directly depends on the amplitude of the light-induced pmf across the thylakoid membrane, we plotted gH^+^ as a function of ECS_T_ (**Figure 4C**). Indeed, we observed that high ECS_T_ values correlated with a partial inhibition of ATP synthase activity. When ATP synthase activity at similar ECS_T_ values were compared, no differences between the wild types and the *pgr5* mutants could be observed anymore, indicating that in the *pgr5* mutants, ATP synthase activity as such is not regulated differently from the wild type. Here, the additional light intensity of 193 µE m^-2^ s^-1^ for Col-0 was used to exactly match the ECS_T_ value, at which gH^+^ of the *pgr5* mutants peaked. On the other hand, despite their higher ECS_T_ values at each tested light intensity, when gH^+^ was plotted against ECS_T_, the two *cgl160* mutants again showed a down-shifted gH^+^ at comparable ECS_T_ values, in line with their lower ATP synthase contents.

### Despite full restoration of thylakoid membrane energization, the electron transport chain of *cgl160 pgr5* is still over-reduced

Usually, the rate-limiting step of linear electron transport is the re-oxidation of plastoquinol by the *b_6_f* (reviewed by Anderson, 1992; Schöttler and Toth, 2014). In consequence, both the acceptor side of PSII and the plastoquinone pool, which are oxidized in darkness, get progressively reduced with increasing light intensity, while the redox-active components of the high-potential chain between cytochrome f and P_700_, which are reduced in darkness, get increasingly oxidized, because electron transfer from plastoquinol via the *b_6_f* is limiting.

When we analyzed the effect of the *cgl160 pgr5* double mutants on the redox state of the different components of the electron transport chain, surprisingly, even though the pmf across the thylakoid membrane and NPQ were fully restored, the electron transport chain remained far more reduced than in the wild types, behaving very similar to the two *pgr5* single mutants (**Figure 5**). According to the chlorophyll-*a* fluorescence parameter qL, the PSII acceptor side was more reduced under light-limited conditions in the *pgr5* mutant (**Figure 5A**). So far, this observation was mainly explained with the compromised NPQ in these mutants. In *cgl160*, due to the higher pmf and “photosynthetic control” of plastoquinol re-oxidation, the PSII acceptor side was more reduced under light-limited conditions as well (Fristedt et al., 2015). However, this effect was much less pronounced than the shift in the light response curve of qL in the *pgr5* mutants. Remarkably, all four double mutants behaved similar to the two single *pgr5* mutants, even though their NPQ was fully restored (**Figure 3B**). Therefore, the massive reduction of the PSII acceptor side in the double mutants cannot be attributed to impaired dissipation of excess excitation energy via NPQ.

**Figure 5:**
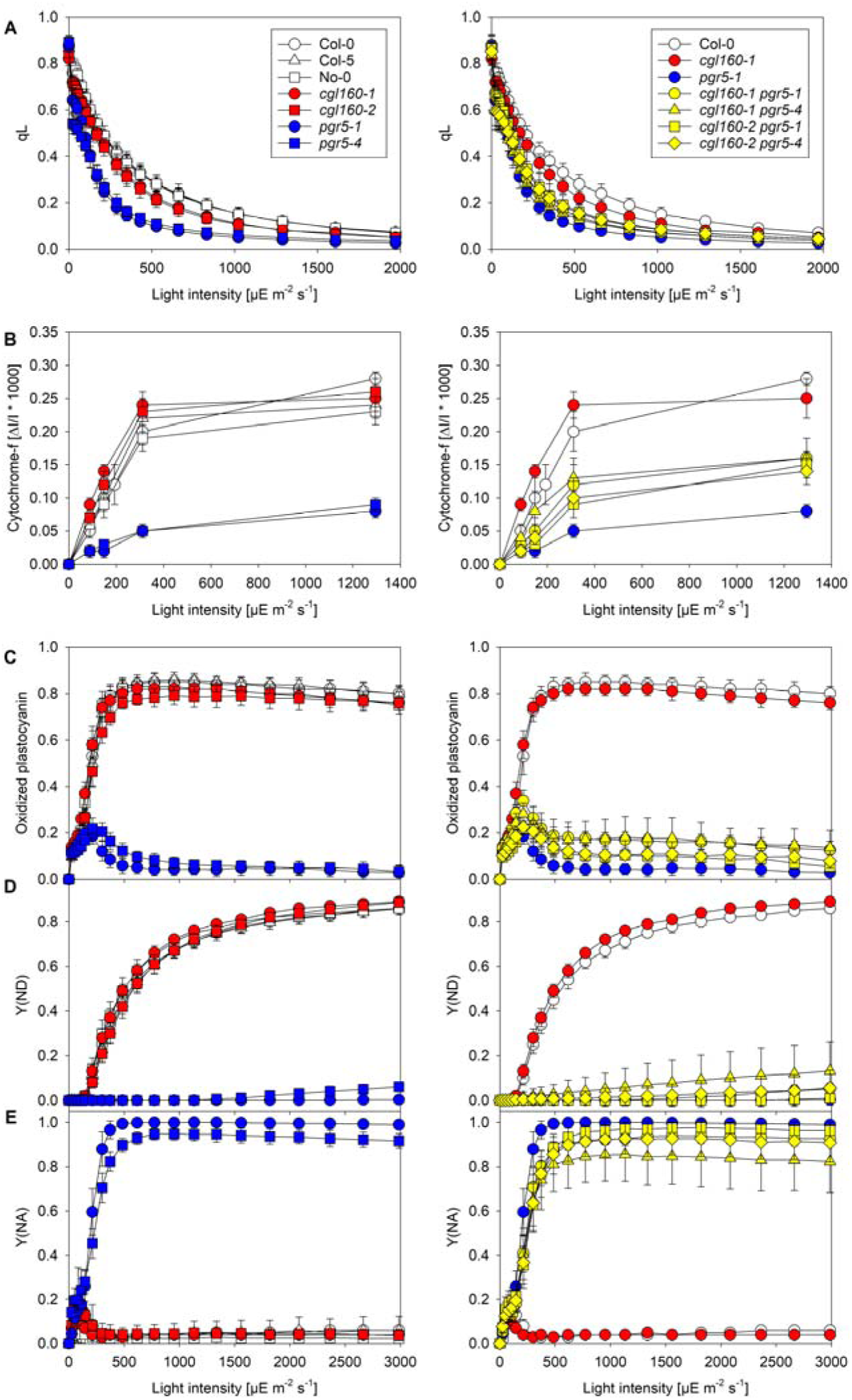
Steady-state redox states of the electron transport chain. Light response curves of qL (A), redox states of cytochrome f (B) and plastocyanin (C), donor-side limitation Y(ND) of PSI (D), and acceptor-side limitation Y(NA) of PSI (E). Left column: Data for the three wild-type accessions and the two single mutants of *cgl160* and *pgr5*, respectively. Right column: Data for Col-0, *cgl160-1*, *pgr5-1* and for the four *cgl160 pgr5* mutants. Please note that because the measurements were done with different instruments optimized for the detection of each individual signal, actinic light intensities in the light response curves differ.

Next, the steady-state redox state of cytochrome f was evaluated. Here, amplitudes of the difference transmittance signals per leaf area are shown (**Figure 5B**). These were measured in parallel to the ECS_T_ and gH^+^ measurements and are the amplitude between the (partly) oxidized state of cytochrome f in the light and its fully reduced state, which was reached within 500 ms after the interruption of actinic illumination. Because the absolute quantifications of the *b_6_f* (**Table 1**) revealed similar complex contents per leaf area in the wild type and all mutants, also similar maximum amplitudes of the light-induced cytochrome f difference transmittance signal would be expected. However, in both *pgr5* mutants, the maximum amplitude of the difference transmittance signal of cytochrome f was reduced by more than 65%, relative to the respective wild types. This shows that in *pgr5,* a large fraction of cytochrome f cannot get oxidized. On the other hand, in the two *cgl160* mutants, under light-limited conditions, cytochrome f was more oxidized than in the wild type, due to the activation of photosynthetic control of plastoquinol oxidation already under light-limited conditions (Correa Galvis et al., 2020). The maximum amplitude of the difference transmittance signal was indistinguishable between the *cgl160* mutants and the wild type. Remarkably, similar to the *pgr5* mutants, the *cgl160 pgr5* mutants showed lower light-induced difference transmittance signal amplitudes than expected from the absolute quantification of the *b_6_f*. Compared to the *pgr5* single mutants, the double mutants showed a somewhat more oxidized redox state of cytochrome f throughout the entire light response curve, with up to 50% of total cytochrome f oxidized in saturating light.

To assess steady-state redox states of plastocyanin (**Figure 5C**) and P_700_ (**Figure 5D**), both were fully oxidized by selective excitation of PSI with far-red light, followed by a short saturating light pulse. The amplitude between the fully reduced state in darkness after the end of the saturation pulse, and the fully oxidized state right at the beginning of the saturating light pulse, was used as a measure for the total amount of redox-active plastocyanin and P_700_, respectively. Based on the known maximum difference signal, for any given light intensity, the relative proportion of oxidized to total plastocyanin and P_700_ was determined (Schreiber and Klughammer, 2016). Similar to the situation with cytochrome f, in the wild type and in both *cgl160* mutants, with increasing light intensity, both plastocyanin (**Figure 5C**) and P_700_ (**Figure 5D**) became oxidized. However, in *pgr5*, a massive impairment of the steady-state oxidation of plastocyanin and P_700_ was observed. Even at the highest actinic light intensity, both remained largely in their reduced state, as previously reported for *pgr5* and *pgrl1* by Takagi and Miyake (2018). In the *cgl160 pgr5* double mutants, the Y(ND) was basically unaltered, relative to the *pgr5* single mutants.

Taken together, these data suggest that processes downstream of PSI, which consume electrons provided by the light reactions, are causal for the poor oxidation of the high-potential chain in all mutants with the *pgr5* genetic background. This is further confirmed by the massive acceptor-side limitation Y(NA) determined for both *pgr5* single mutants and the four *cgl160 pgr5* double mutants (**Figure 5E**). While in the wild types and the *cgl160* mutants, Y(NA) only transiently occurred under light-limited conditions, which is attributable to an incomplete activation of the CBC and downstream reactions of primary metabolism, in the *pgr5* single and the *cgl160 pgr5* double mutants, Y(NA) increased with light intensity, suggesting a permanent congestion of electron consuming processes of primary metabolism.

## Discussion

The *pgr5* mutant is characterized by impaired thylakoid membrane energization (**Figure 4A**), impaired NPQ (**Figure 3B**), and an over-reduction of the PSI acceptor side (**Figure 5E**). These defects have been explained by at least four fundamentally different scenarios: In the original **scenario 1**, all defects were attributed to impaired CEF, leading to reduced pmf formation across the thylakoid membrane (Munekage et al., 2002). The reduced pmf was then assumed to be causal for the over-reduction of the PSI acceptor side, due to impaired NPQ and absence of photosynthetic control of plastoquinol re-oxidation at the *b_6_f* (Munekage et al., 2002; Zhou et al., 2023). In this scenario, the abolished massive acceptor side limitation of PSI after the introduction of FLVs into *pgr5* was interpreted as mediated by the restoration of pmf formation and photosynthetic control (Yamamoto et al., 2016; Shikanai and Yamamoto, 2017; Yamamoto et al., 2020).

In **scenario 2**, mainly based on the observation by Avenson et al. (2005) that feedback inhibition of ATP synthase was impaired in pgr5, Tikkanen et al. (2015) and Rantala et al. (2020) speculated that PGR5 exerts feedback-regulation on photosynthetic light reactions by fine-tuning ATP synthase activity. In **scenario 3**, an ATP deficit due to a slight decrease in CEF was assumed to result in elevated levels of ADP and Pi, thereby resulting in deregulated chloroplast ATP synthase activity, which in turn resulted in the massively reduced pmf formation. In this scenario, the over-reduction of the PSI acceptor side was explained by the metabolic disturbance due to limited ATP provision, which would then also restrict metabolic consumption of NADPH (Avenson et al., 2005). A **fourth scenario** interpreted the phenotypes in *pgr5* as the consequence of a restriction in electron transport processes downstream of PSI, possibly due to inefficient recruitment of ferredoxin to the thylakoid membrane and impaired ferredoxin reduction. Then, this congestion in linear electron transport was suggested to cause the over-reduction of the electron transport chain (Takagi and Miyake, 2018). Likewise, in *C. reinhardtii*, a role of PGR5 and PGRL1 in FNR binding to a PSI supercomplex was suggested (Mosebach et al., 2017).

Here, we disprove scenario one. We crossed two independent *pgr5* mutants with two mutants of the auxiliary protein CGL160, which supports ATP synthase assembly (Rühle et al., 2014). Loss of CGL160 results in a massive reduction in ATP synthase accumulation and decreased ATP synthase activity (Fristedt et al., 2015; Correa Galvis et al., 2020). In the *cgl160 pgr5* double mutants, pmf formation was fully restored, as shown by light response curves of ECS_T_ (**Figure 4A**). As a consequence, also NPQ in the double mutants was fully restored (**Figure 3B**). These results are similar to the introduction of FLVs into *pgr5*, which also restored pmf formation (Yamamoto et al., 2016; Shikanai and Yamamoto, 2017; Yamamoto et al., 2020). However, different to the situation after introduction of FLVs into *pgr5*, the *cgl160 pgr5* double mutants still suffer from a massive over-reduction of the entire electron transport chain, from the PSII acceptor side (qL; **Figure 5A**), through cytochrome f (**Figure 5B**) and plastocyanin (**Figure 5C**) to P_700_ (**Figure 5D**). This is similar to the *pgr5* single mutants, and suggests that the original defect in *pgr5* is related to the massive limitation downstream of the PSI acceptor side (**Figure 5E**), in agreement with both scenario 3 and scenario 4. Defects in both NPQ and photosynthetic control are a secondary consequence of the impaired electron distribution downstream of PSI. Due to this congestion, the high potential chain remains reduced even in saturating light, which in turn limits *b_6_f* activity and pmf formation by the Q cycle. If this interpretation of events is correct, the introduction of FLVs would directly abolish the impaired electron distribution downstream of PSI by providing an additional major electron sink, instead of mainly acting via the re-establishment of both pmf and photosynthetic control (Yamamoto et al., 2016; Shikanai and Yamamoto, 2017; Yamamoto et al., 2020). Likewise, any reduction in linear electron transport would reduce the congestion downstream of PSI and thereby alleviate the phenotype of the *pgr5* mutant. An example for this is the sixtuple mutant of *pgr5* crossed with the Δ5 mutant combining knockout mutations in the OEC subunit genes *PSBO1*, *PSBP2*, PSBQ1, *PSBQ2*, and *PSBR*, with massively reduced PSII activity (Suorsa et al., 2016).

### PGR5 does not directly affect ATP synthase activity

Even though wild-type plants and *pgr5* mutants contain the same amount of chloroplast ATP synthase (**Figure 2**), Avenson et al. (2005) observed increased ATP synthase activity in *pgr5*. These differences became most pronounced when experiments were performed under CO_2_-limited conditions (50 ppm CO_2_), which resulted in repressed ATP synthase activity in the wild type. Based on these data, Tikkanen et al. (2015) and Rantala et al. (2020) speculated in scenario 2 that PGR5 senses the redox state of PSI electron acceptors and exerts feedback-regulation on photosynthetic light reactions by fine-tuning ATP synthase activity via an unknown mechanism.

Our data clearly argue against a specific defect in ATP synthase regulation in the absence of PGR5, and are in full agreement with the original explanation of the increased ATP synthase activity by Avenson et al. (2005). When wild-type plants were illuminated with a low light intensity, to exactly match the light-induced pmf across the thylakoid membrane to that in the *pgr5* mutants under light saturation, their ATP synthase activities were indistinguishable from the *pgr5* mutants (**Figure 4A to C**). The simplest explanation for the partial inactivation of ATP synthase at high pmf values is that in the wild type in saturating light, photosynthesis is limited either by the CBC or by downstream processes such as triose-phosphate utilization for starch and sucrose biosynthesis (Sharkey and Vanderveer, 1989). Then, metabolic regeneration of Pi restricts ATP synthase activity (Kanazawa and Kramer 2002; Kiirats et al. 2009; McClain et al., 2023), in turn leading to the increased pmf across the thylakoid membrane. Only at the lower pmf values, when ATP synthase activity is highest, no feedback inhibition of ATP synthesis by downstream metabolic limitations occurs. It is noteworthy that when Yamamoto and Shikanai (2020) measured gH^+^ at different light intensities in wild type and *pgr5*, they did not observe an increase in wild-type activity under low light conditions, and wild-type activity was always lower than that of *pgr5*, seemingly contradictive to our results. However, they determined ATP synthase activity on dark-adapted detached leaves with a very short pre-illumination time prior to the measurements, suggesting that the CBC was not fully activated, and therefore limited the regeneration of ADP and Pi for ATP synthase activity even under light-limited conditions.

### *b_6_f* activity is strongly repressed in the *cgl160 pgr5* mutants

So far, we have demonstrated that neither a de-regulation of chloroplast ATP synthase, nor the impaired thylakoid membrane energization are directly causal for the over-reduction of both high potential chain and of the PSI acceptor side. However, we have not considered whether photosynthetic control worked normally in the *cgl160 pgr5* double mutants. The typical effects of photosynthetic control, a slower oxidation of plastoquinol and slower reduction of plastocyanin and P_700_ during dark-interval kinetics after saturating illumination (for example: Rott et al., 2011), cannot be determined when a general over-reduction of the electron transport chain occurs and the fully oxidized states of the components of the high potential chain cannot be reached. Therefore, we instead looked at the enzymatic turnover number of the *b_6_f* (Table 1), which was calculated from the capacity of linear electron transport per leaf area (**Figure 3A**) and the content of the *b_6_f* per leaf area (**Table 1**). For the three wild types, average turnover numbers of more than 260 electrons s^-1^ were calculated, which is close to the maximum enzymatic turnover number of 250 to 300 electrons s^-*1*^ reported for the *b_6_f* from *C. reinhardtii* (Pierre et al., 1995) and tobacco (Hojka et al., 2014). All mutants showed a reduced enzymatic activity of the *b_6_f*. In case of the *cgl160* mutants with turnover numbers of around 200 s^-1^, this is directly attributable to their increased pmf and induction of photosynthetic control (Fristedt et al., 2015; Correa Galvis et al., 2020). Also in *pgr5*, the turn-over number of the *b_6_f* was slightly decreased, which could be explained by linear electron transport being slowed-down by the limitation downstream of PSI, in accordance with both scenario 3 and scenario 4. Remarkably, the turnover number of the *b_6_f* was by far lowest in the *cgl160 pgr5* double mutants. This cannot be explained by the downstream limitation of electron transport alone, because the steady-state oxidation level of cytochrome f was slightly increased, relative to the *pgr5* single mutants (**Figure 5B**). A closer inspection of both thylakoid membrane energization **(Figure 4A**) and NPQ (**Figure 3B**) showed that due to repressed ATP synthase accumulation, the *cgl160 pgr5* double mutants have a slightly higher light-induced pmf across the thylakoid membrane already under light limited conditions than the wild type, similar to the *cgl160* mutants. Likely, the limitation in electron transport in the high potential chain by processes downstream of PSI occurring in *pgr5* (**Figure 3A**), and the inhibition of linear electron transport in *cgl160* via the induction of photosynthetic control, have an additive negative effect on *b_6_f* activity in *cgl160 pgr5*, which is co-limited by plastoquinol re-oxidation due to photosynthetic control, and a lower oxidizing power provided by the high-potential chain. Ultimately, this may also explain why growth of the double mutants is strongly retarded, and biomass accumulation is impaired (**Figure 1**).

### What is the molecular function of PGR5?

Our data clearly rule out scenarios one and two as explanation of the *pgr5* phenotype. However, the *cgl160 pgr5* double mutants do not allow us to clearly distinguish between scenario 3, where a minor reduction in CEF leads to a perturbed metabolic consumption of reducing equivalents, and therefore results in the congestion of the electron transport chain, and scenario four, which assumes a direct limitation in electron transport processes downstream of PSI, possibly at the level of ferredoxin (Takagi and Miyake, 2018) or FNR (Mosebach et al., 2017). If according to scenario 3, a deficit in ATP production due to impaired CEF is causal for the congestion downstream of PSI, similar defects should be observed in a large number of metabolic situations leading to an increased metabolic demand for ATP, relative to NADPH. For example, high rates of starch biosynthesis or especially photorespiration massively increase the demand for ATP, while other major pathways such as nitrite reduction have a higher demand for reducing equivalents (discussed in detail by: Noctor and Foyer, 1998; Noctor and Foyer, 2000; Fu and Walker, 2023). However, to our knowledge, neither under conditions of high photorespiration, nor under other conditions selectively increasing ATP demand, defects similar to those in *pgr5* have been observed. Furthermore, chloroplast evolved multiple flexibility mechanisms to cope with fluctuating ATP demands. In addition to the different pathways of CEF, these flexibility mechanisms include the export of excess reducing equivalents from the chloroplast via the malate valve (Scheibe, 2004; Hebbelmann et al., 2012) and other metabolic shuttles such as the export of triose-phosphate to and import of 3-phosphoglycerate from the cytosol (Noctor and Foyer, 2000). Alternatively, excess electrons can be directly transferred to O_2_ via the PTOX (Bolte et al., 2020) or the Mehler-Asada cycle. Most of these pathways have a relatively low activity in most C_3_ plants, but when the function of one pathway is disturbed, the other flexibility mechanisms are upregulated. Interestingly, in the *pgr5* mutant, export of excess reducing equivalents from the chloroplast to the mitochondria might play a major role, because the initial activity of both the NADPH-dependent malate dehydrogenase as key enzyme of the malate valve and of the mitochondrial alternative oxidase (AOX) are strongly increased in *pgr5* under high light (Yoshida et al., 2007). When the AOX is inactivated, an *aox1a pgr5-1* double mutant suffers from clearly impaired growth, relative to both *pgr5* and *aox1a*, both under low – and high-light conditions (Yoshida et al., 2011). These results are compatible both with scenario three and four, and strongly agree with a limitation in metabolic NADPH consumption in the chloroplast, which then activates the malate valve.

In support of scenario four, the NDH pathway is believed to play the major role in covering an ATP deficit in the chloroplast, because it has a proton-pumping activity itself, so that the NDH pathway can generate a much higher pmf per electron than the FQR pathway (Strand et al., 2017). Also, the FQR pathway is strongly inhibited already at relatively low ATP concentration, while the NDH complex pathway is much less sensitive to feedback inhibition by ATP, again suggesting that it is the main pathway to compensate for an ATP depletion (Fisher et al., 2019). Therefore, the defects in *pgr5* are unlikely to be caused by an impaired function of the FQR pathway of CEF and a congestion of the electron transport chain occurring in response to limitations in NADPH consumption because ATP supply is low.

If a restriction in ATP provision by CEF is not causing the limited metabolic consumption of electrons downstream of PSI, what else might be the primary defect in *pgr5*? To assess a limitation directly downstream of PSI in the *pgr5* and *cgl160 pgr5* mutants ourselves, we performed ferredoxin measurements in our mutants, similar to Takagi and Miyake (2018). However, we observed major complications due to electron accumulation on the 4Fe4S clusters on the PSI acceptor side in both single and double mutants, which resulted in difference transmittance changes very similar to those of ferredoxin. Due to these signal contaminations, we decided against including our ferredoxin data in this manuscript. Even though we therefore could not directly confirm the impaired ferredoxin reduction suggested by Takagi and Miyake (2018), we agree that very likely, the main problem in the *pgr5* mutant is a defect in electron partitioning downstream of PSI, in line with scenario 4. If this problem occurs on the level of ferredoxin, the FNR (Mosebach et al., 2017), their recruitment to the thylakoid membrane, or even further downstream still remains to be determined. A direct defect in forward electron transfer to ferredoxin and NADPH might be difficult to reconcile with the full restoration of linear electron transport by the introduction of FLVs (Yamamoto et al., 2016; Shikanai and Yamamoto, 2017; Yamamoto and Shikanai, 2020), because FLVs accept their electrons from NADPH. Therefore, a defect in electron transfer between PSI and NADPH should not be rescued by FLVs. A defect in electron transfer prior to NADPH reduction is also incompatible with the increased activation of the NADPH-dependent malate dehydrogenase in *pgr5*, which also requires NADPH as electron donor (Yoshida et al., 2007).

Using a bioluminescence-based ATP indicator, a slower increase in ATP concentration upon illumination was recently shown in *pgr5* (Sato et al., 2018). However, because no parallel assessment of changes in NADPH levels was done, it is not possible to conclude on the primary limiting factor for photosynthetic electron transport and metabolism from these data. Recently, in *C. reinhardtii*, a severe restriction in starch biosynthesis was shown to cause an acceptor side limitation of PSI and result in over-reduction of the entire electron transport chain, from the PSII acceptor side to restricted light-induced oxidation of cytochrome f (Saroussi et al., 2023). These defects closely resemble those in *pgr5*, demonstrating that also a defect in the reactions of primary chloroplast metabolism might be causal for the defects observed in *pgr5*. To ultimately identify the precise molecular function of PGR5, it might be necessary to determine the impact of the *pgr5* mutant on adenylate energy charge, Pi levels, and redox poise in the chloroplasts. Such parallel measurements would provide indications if a major imbalance between ATP and NADPH provision, and therefore a PGR5 function somehow related to CEF as described in scenario 3, could explain the observed defects. Furthermore, by measurements of flux rates, enzyme contents and enzyme activation states of the major metabolic pathways in the chloroplast, a possible role of PGR5 in controlling processes with a high demand for reducing energy, which could explain the electron congestion of the photosynthetic apparatus, could be addressed. Besides the CBC itself, also reactions such as nitrite reduction and fixation in organic molecules might be of interest. Such studies should ultimately allow the identification of the precise molecular function of PGR5 in chloroplast bioenergetics and redox poising.

## Methods

### Identification of the new *pgr5-4* mutant

For structural annotation of the *pgr5-4 Ds*– transposon insertion, Illumina DNA sequencing reads were mapped against the *A. thaliana* reference genome (TAIR10, www.arabidopsis.org) using bwa mem (Li, 2013). The mapping files were processed using SAMtools (Li et al., 2009). The exact insertion site was identified by visualizing mappings in IGV (Thorvaldsdóttir et al., 2013)). The inserted *Ds*-transposon sequence was reconstructed using edge extension assembly from both sides of the insertion site. Reads not mapped to the reference genome were parsed for exact matches with bordering sequences. They were then multiple sequence aligned using MUSCLE (Edgar, 2004) to extract a consensus sequences to be appended. This process was reiterated using the appended border sequences until full reconstruction. The final sequence model was checked by mapping Illumina sequencing reads against the *A. thaliana* reference plus the reconstructed model and visually checking for consistency in IGV.

### Plant cultivation

To synchronize germination, wild-type Arabidopsis accessions Col-0, Col-5 and No-0 and transgenic seeds were stratified for 2 d at 4°C in 1% agarose with 0.3% gibberellic acid A4 and A7 (0.2% and 0.1% [v/v], respectively). Seeds were sown on soil, grown in long-day conditions (16-h light/8-h darkness at 20°C/ 6°C, 75%/75% relative humidity) at 250 µE m^-2^ s^-1^ for 7 d. Plants were then moved to long-day conditions with 150 µE m^-2^ s^-1^, 16°C night temperature and 60% day relative humidity. After 2 weeks, plants were pricked into individual pots. All measurements were performed with plants that were 4 to 6 weeks old.

### Thylakoid membrane isolation and photosynthetic complex quantification

For thylakoid isolations, plants were kept in darkness for 24 hours before rosette leaves from five to eight individual plants were harvested and pooled. Then, thylakoid membranes were isolated according to Schöttler et al. (2004). Chlorophyll content and a/b ratio were determined in 80% (v/v) acetone according to Porra et al. (1989). The contents of PSII and the *b_6_f* were determined from difference absorbance signals of cytochrome b_559_ (PSII) and the cytochromes b_6_ and f in destacked thylakoids equivalent to 50 µg chlorophyll mL^−1^ (Kirchhoff et al., 2002). All cytochromes were fully oxidized by the addition of 1 mM potassium hexacyanoferrate (III), and stepwise reduced by the addition of 10 mM sodium ascorbate to reduce the high-potential form of cytochrome b_559_ and cytochrome f, and the addition of 10 mM sodium dithionite to reduce the low potential form of cytochrome b_559_ and the two b-type hemes of cytochrome b_6_. Using a V-650 spectrophotometer equipped with a head-on photomultiplier (Jasco GmbH, Germany), at each of the three redox potentials, absorbance spectra were measured between 575 and 540 nm wavelength. The spectral bandwidth was 1 nm and the scanning speed 100 nm min^-1^. Ten spectra were averaged per redox condition. Difference spectra were calculated by subtracting the spectrum measured in the presence of hexacyanoferrate from the ascorbate spectrum, and by subtracting the ascorbate spectrum from the spectrum measured in the presence of dithionite, respectively. Finally, a baseline calculated at wavelengths between 540 and 575 nm was subtracted from the signals. Then, the difference spectra were deconvoluted using reference spectra as previously described (Kirchhoff et al., 2002; Lamkemeyer et al., 2006).

PSI was quantified from light-induced difference absorbance changes of P_700_. Thylakoids equivalent to 50 µg chlorophyll mL−1 were solubilized in the presence of 0.2% (w/v) n-dodecyl-β-D-maltoside (DDM). After the addition of 10 mM sodium ascorbate as the electron donor and 100 µM methylviologen as the electron acceptor, P_700_ was photo-oxidized by a 250 ms light pulse of (2000 µmol photons m^−2^ s^−1^). Measurements were performed with the plastocyanin-P700 version of the Dual-PAM instrument (Heinz Walz GmbH, Effeltrich, Germany). Plastocyanin contents, relative to PSI, were determined in intact leaves (see below) and then recalculated based on the absolute PSI quantification in isolated thylakoids.

### Protein gel electrophoresis and immunoblotting

Thylakoid proteins were separated by SDS-PAGE and then transferred to a polyvinylidene membrane (Hybond P, GE Healthcare) using a tank blot system (Perfect Blue Web M, VWR International GmbH, Darmstadt, Germany). Immunochemical detection was performed using an enhanced chemiluminescence detection reagent (ECL Prime, GE Healthcare) according to the manufacturer’s instructions. Chemiluminescence was detected using the G:BOX XT4 imaging system (Syngene, Cambridge, UK). The following specific antibodies against the photosynthetic proteins were purchased from Agrisera AB (Vennäs, Sveden): PsbC (order number: AS11 1787), PsbD (AS06 146), PSBO (AS05 092) PetA (AS06 119), PetB (AS03 034), PETC (AS08 330), PsaA (AS06 172), PSAL (AS06 108), AtpB (AS05 085) and ATPC (AS08 312).

### Chlorophyll-*a* fluorescence measurements

*In vivo* parameters of chlorophyll-*a* fluorescence were determined with the fiberoptics version of the DUAL PAM-100 instrument (Heinz Walz GmbH, Effeltrich, Germany). Light-response curves of linear electron flux, non-photochemical quenching (NPQ; Baker et al., 2007), and the redox state of the PSII acceptor side (qL; Kramer et al., 2004) were measured after 30 min of dark adaptation. The light intensity was increased stepwise from 0 to 2000 µE m^−2^ s^−1^, with a measuring time of 180 s per light intensity under light-limited conditions and of 60 s under light-saturated conditions.

### ATP synthase activity and light-induced pmf

The thylakoid membrane conductivity for protons (gH+) was used as a measure for ATP synthase activity (Baker et al., 2007). It was determined on intact leaves from the decay kinetics of the electrochromic shift signal (ECS) during a short interval of darkness. The experiments were performed at 22°C. Plants were directly taken from growth chambers, and leaves were then pre-illuminated for 5 min with saturating light (1295 µE m^−2^ s^−1^) so that photosynthesis was fully activated and ATP synthase activity was not limited by ATP consumption by the CBC. The saturating illumination was interrupted by 10 s intervals of darkness, and 10 repetitions of the dark interval relaxation kinetic were measured and averaged, to improve the signal to noise ratio. Then, the actinic light intensity was step-wise decreased to 311, 193 (only Col-0), 148 and 88 µE m^-2^ s^-1^. At each light intensity, plants were again pre-illuminated for 5 min, to allow photosynthesis to fully adapt to the new light intensity, and dark interval relaxation kinetics were averaged. Finally, the light intensity was again increased to 1295 µE m^−2^ s^−1^, to ensure that the measurement procedure had not altered ATP synthase activity.

To determine gH^+^, the rapid first phase of the decay kinetic of the electrochromic shift during the first 120 to 250 ms of darkness (depending on ATP synthase activity at the respective light intensity and in the respective mutant, due to the slower decay in all plants with the *cgl160* mutant background) was fitted with a single exponential decay function. The reciprocal value of the time constant was used as a measure of ATP synthase activity. Signals were measured and deconvoluted using a KLAS-100 spectrophotometer (Heinz Walz GmbH, Effeltrich, Germany) as previously described (Rott et al., 2011). The maximum amplitude of the ECS during the first phase of its relaxation kinetic was also used as a measure for the total light-induced pmf across the thylakoid membrane (ECS_T_). To take differences in leaf chlorophyll content into account, the signal amplitude was normalized to the chlorophyll content per leaf area (Rott et al., 2011).

### Spectroscopic measurements of cytochrome f redox kinetics

Leaves were pre-illuminated as described for the ATP synthase measurements to fully activate the CBC and – if possible – avoid an acceptor-side limitation of PSI. Cytochrome f was fully oxidized by a saturating light pulse, followed immediately by a short interval of darkness, to allow its complete reduction. The fully reduced state of cytochrome f was reached after a maximum of 500 ms in darkness, and the amplitude of the difference transmission signal between the fully oxidized state during the light pulse and the fully reduced state in darkness was used as a measure of redox-active cytochrome f. However, as demonstrated by major discrepancies between the light-induced difference absorbance signals of cytochrome f and the maximum content of the *b6f* determined in vitro (see results), in all mutants containing a *prgr5* background, it was impossible to achieve a full oxidation level of cytochrome f. The cytochrome f signal was measured with the KLAS-100 spectrophotometer and deconvoluted as previously described (Klughammer et al. 1990; Rott et al. 2011).

### Steady-state redox states of plastocyanin and P700

PSI-related measurements were performed with the plastocyanin-P_700_ version of the Dual-PAM instrument (Schöttler et al., 2007). Plants taken directly from the growth chamber were pre-illuminated with saturating light for 3 min prior to measurements, to activate CBC and avoid an acceptor-side limitation of PSI. After 10 s in the dark, the maximum difference absorbance signals of redox-active plastocyanin and PSI were quantified by far-red illumination for eight seconds, followed by a short saturating light pulse. The signal amplitude between the fully reduced state prior to illumination and in darkness after the end of the saturating light pulse, and the fully oxidized state of both plastocyanin and P700 at the beginning of the saturating light pulse, was used as a measure for the total amount of redox-active plastocyanin and P_700_, respectively. After determination of the maximum amplitude, the actinic light intensity was stepwise increased as described above for the chlorophyll-a fluorescence measurements. The fraction of PSI reaction centers limited at the donor side, Y(ND), and on the acceptor side, Y(NA), was determined as described (Schreiber and Klughammer, 2016).

## Author Contributions

UA and MAS designed the project. SK generated and genotyped the mutants and performed growth analyses. SGT and AH isolated thylakoid membranes and quantified photosynthetic complexes. WT performed immunoblots. MAS and WT performed in vivo photosynthetic measurements including chlorophyll-a fluorescence and difference absorbance measurements. MAS wrote the article with input from all co-authors. All authors agreed to the publication of this manuscript.

## Acknowledgements

The authors thank the Max-Planck society for providing funding for the project. They also thank the MPI-MP Green Team for plant care and cultivation.

